# Salicylic acid perturbs sRNA-gibberellin regulatory network in immune response of potato to *Potato virus Y* infection

**DOI:** 10.1101/130757

**Authors:** Maja Križnik, Marko Petek, David Dobnik, Živa Ramšak, Špela Baebler, Stephan Pollmann, Jan Kreuze, Jana Žel, Kristina Gruden

## Abstract

*Potato virus Y* is the most economically important potato viral pathogen. We aimed at unraveling the roles of small RNAs (sRNAs) in the complex immune signaling network controlling the establishment of tolerant response of potato cv. Désirée to the virus. We constructed a sRNA regulatory network connecting sRNAs and their targets to link sRNA level responses to physiological processes. We discovered an interesting novel sRNAs-gibberellin regulatory circuit being activated as early as 3 days post inoculation before viral multiplication can be detected. Increased levels of miR167 and phasiRNA931 were reflected in decreased levels of transcripts involved in gibberellin biosynthesis. Moreover, decreased concentration of gibberellin confirmed this regulation. The functional relation between lower activity of gibberellin signaling and reduced disease severity was previously confirmed in Arabidopsis-virus interaction using knockout mutants. We further showed that this regulation is salicylic acid-dependent as the response of sRNA network was attenuated in salicylic acid-depleted transgenic counterpart NahG-Désirée expressing severe disease symptoms. Besides downregulation of gibberellin signaling, regulation of several other parts of sRNA network in tolerant Désirée revealed similarities to responses observed in mutualistic symbiotic interactions. The intertwining of different regulatory networks revealed shows how developmental signaling, disease symptom development and stress signaling can be balanced.

## Introduction

Potato (*Solanum tuberosum* L.) is the world’s most important non-grain staple crop. Viruses pose a serious threat to potato production, not only because of the effects caused by the primary infection but also because potato is propagated vegetatively so that viruses are transmitted through the tubers and accumulate over time (Solomon-Blackburn and Barker, 2001). The most devastating potato virus is *Potato virus Y* (PVY) (Karasev and Gray, 2013). PVY is a member of the Potyviridae family and comprises of many diverse strains. Worldwide, the most harmful strain is PVY^NTN^ which has been responsible for huge decreases in quality and quantity of potato tuber production (Scholthof et al., 2011). One of the most widely grown potato cultivars is cv. Désirée, which is tolerant to PVY^NTN^, meaning that the virus replicates and spreads systemically, however, symptoms of the disease are mild or not visible at all (Baebler et al., 2011; Kogovšek and Ravnikar, 2003). Tolerance may have an advantage over resistance for crop protection because it does not actively prevent virus infection and/or replication, therefore there is little evolutionary pressure for PVY to mutate and to evolve into more aggressive strains (Bosch et al., 2006). Hence, the tolerant phenotype is likely to be more durable than resistance (Wilson, 2014). Until now, studies on Désirée-PVY^NTN^ interactions have focused on the detection of changes in the plant transcriptome and proteome, particularly those related to plant hormonal signaling (Baebler et al., 2011; Stare et al., 2015) where salicylic acid (SA) was found to be the crucial component for attenuation of the disease symptoms (Baebler et al., 2011). Nevertheless, understanding of the mechanisms that underlie tolerance response to the virus is still incomplete.

RNA silencing is a basal antiviral system in plants, where DICER-like (DCL) proteins cleave viral dsRNA structures, giving rise to virus-derived small interfering RNAs (vsiRNAs), which are then incorporated into Argonaute (AGO) protein(s) to guide viral RNA degradation (Baulcombe, 2004). To counter this host defence mechanism, viruses have evolved viral suppressors of RNA silencing (Csorba et al., 2015). Helper component-proteinase (HCPro) of potyviruses suppresses silencing by sequestering small RNAs (sRNAs) and AGO1 and thus counteracts the degradation of viral RNA (Ivanov et al., 2016). Another level of plant antiviral defence is mediated by resistance genes, leading towards effector-triggered immunity, often resulting in hypersensitive response and programmed cell death (Coll et al., 2011; Zvereva and Pooggin, 2012). The cv. Désirée carries the *Ny* gene conferring resistance against strain PVY^O^, but lacks resistance genes against the PVY^NTN^ strain (Singh et al., 2008) and thus does not respond by triggering an efficient effector-triggered immunity.

Recent findings revealed that endogenous RNA silencing mediated by microRNAs (miRNAs) and small interfering RNAs (siRNAs) could play important roles in plant immunity (Li et al., 2011; Navarro et al., 2005; Seo et al., 2013; Weiberg and Jin, 2015). These 18-24-nt long non-coding sRNAs are able to negatively regulate gene expression by binding to the specific mRNA targets which leads to either promoting their degradation, inhibiting their translation, or suppressing transcription by epigenetic modification (Baulcombe, 2004). The endogenous RNA silencing can be amplified by the production of secondary phased siRNAs (phasiRNAs), triggered by 22-nt miRNAs/siRNAs (Chen et al., 2010; Cuperus et al., 2010). phasiRNAs are generated in phase relative to positions of the miRNA cleavage site, can be produced from both coding or non-coding transcript (*PHAS* loci) and are able to target transcripts not only in *trans* but also their *PHAS* loci of origin in *cis* and thus additionally contribute to the autoregulation (Borges and Martienssen, 2015). miRNAs have been associated with defence responses against several pathogens (Peláez and Sanchez, 2013; Ruiz-Ferrer and Voinnet, 2009). *Arabidopsis thaliana* miR393 was the first plant miRNA reported to play a key role in antibacterial immunity by repressing auxin signaling (Navarro et al., 2005). Several studies have uncovered the miRNA-mediated silencing of immune receptor gene (*R*-gene) transcripts. Infection by pathogens e.g. viruses or bacteria, relieves the silencing, leading to the accumulation of *R* proteins and activation of immune responses (Li et al., 2011; Park and Shin, 2015; Shivaprasad et al., 2012).

This growing body of evidence suggests that sRNAs are integral components of plant immunity. However, none of the studies performed so far investigated the sRNA regulatory network in potato-virus interaction at the systems level linking it to transcriptional regulation. The aim of this study was to investigate sRNAs’ role in establishment of the tolerant response of potato to PVY^NTN^, hence we have studied response in the early stage of the infection, before the viral multiplication can be detected. Employing high-throughput sequencing technology, we characterized and compared the sRNA expression patterns between PVY-infected and healthy tolerant potato plants. In addition, this information was linked to expression profiles of their target transcripts identified *in silico* and by Degradome-Seq and used for sRNA regulatory network construction. Besides the already described regulation of *R*-gene transcripts, we have discovered a previously undescribed connection between sRNAs and gibberellin (GA) biosynthesis, representing an important link between immune and developmental signaling pathways. This link was confirmed by hormonal content measurements. Additionally, we analyzed sRNA regulatory network in transgenic NahG-Désirée. We showed that response of the discovered sRNA network is attenuated in the absence of SA, indicating a mechanism through which SA is regulating disease tolerance in potato.

## Materials and Methods

### Plant material

Potato leaves of cv. Désirée were mock or PVY^NTN^-inoculated (isolate NIB-NTN, GenBank accession no. AJ585342) as described previously (Stare et al., 2015). Plant material of the inoculated leaves was collected 3 days post inoculation (dpi), corresponding to early stages of viral multiplication. Three and four biological replicates (individual leaves from different plants per group) were analyzed for RNA analysis and for hormonal measurements, respectively. Three plants from each group were monitored for plant height, till 21 dpi, when they all started to senesce. The same experimental set up was designed also for analysis of transgenic NahG-Désirée plants (Halim et al., 2004).

### RNA extraction, library preparation and sRNA sequencing

Total RNA was extracted from 100 mg of homogenized leaf tissue using TRIzol reagent (Thermo Fisher Scientific, Waltham, MA) and MaXtract High Density tubes (Qiagen, Hilden, Germany) following manufacturers’ protocols. RNA concentration, quality and purity were assessed using agarose gel electrophoresis and NanoDrop ND-1000 spectrophotometer (Thermo Scientific). sRNA NGS libraries were generated from total RNA samples using the TailorMix miRNA Sample Preparation Kit (SeqMatic LLC, Fremont, CA) and sequenced on the Illumina HiSeq 2000 Sequencing System at SeqMatic LLC.

### Identification of potato and virus-derived sRNAs

The raw reads were quality filtered using Filter Tool of the UEA sRNA Toolkit (Moxon et al., 2008) by discarding low complexity reads (containing at most two distinct nucleotides), reads shorter than 18 nt and longer than 25 nt, reads matching tRNA/rRNA sequences and reads not mapped to the potato genome (PGSC_DM_v4.3) (Xu et al., 2011). To identify known annotated miRNAs, the remaining reads were compared to all plant miRNAs registered in miRBase database (release 21) (Kozomara and Griffiths-Jones, 2014), allowing no mismatches. The sequences that matched mature miRNAs from other plants than potato (miRNA orthologs), were mapped to the potato genome to find corresponding *MIR* loci able to form hairpin structure (An et al., 2014) and named according to annotation of conserved miRNA (Meyers et al., 2008). miRNAs that had different 5′ and 3′ ends with respect to the mature miRNA, were annotated as isomiRs. To identify novel unannotated miRNAs, filtered reads were submitted to miRCat tool of the UEA sRNA Toolkit using default parameters for plants, considering only reads of lengths 18-24 nt. Reads were first mapped to the potato genome, then the 100 and 200 nt long windows around the aligned reads were extracted (An et al., 2014). The predicted secondary structures were trimmed and analyzed to verify the characteristic hairpin pre-miRNA structure according to plant miRNAs annotation criteria (Meyers et al., 2008). An additional criterion we have imposed was, that novel miRNAs should be present at least in two analyzed samples with more than five raw reads. Potential novel miRNAs were mapped against miRBase and sequences that matched known plant miRNAs with up to three mismatches were excluded. The novelty of potato specific miRNAs was verified with the miRPlant version 5 (An et al., 2014) using default parameters and additionally rechecked against the latest releases of Rfam (Nawrocki et al., 2015); http://rfam.xfam.org/), tRNA (Chan and Lowe, 2016); http://gtrnadb.ucsc.edu/) and snoRNA databases (Yoshihama et al., 2013); http://snoopy.med.miyazaki-u.ac.jp/). Families of novel miRNAs were determined by clustering their sequences with sequences of known miRNAs using CD-HIT with an identity threshold of 0.9 (Huang et al., 2010). To identify vsiRNAs, reads of lengths 20-24 nt from all PVY^NTN^-infected samples were mapped to the reconstructed consensus PVY^NTN^ genome (Kutnjak et al., 2015) using CLC Genomics Workbench version 8 (http://www.clcbio.com/) allowing only 100 % identity.

### Prediction of novel potato phasiRNAs and *PHAS* loci

Prediction of phasiRNAs and phasiRNA-producing loci (*PHAS* loci) was performed using ta-siRNA prediction tool (Chen et al., 2007; Moxon et al., 2008) utilizing the potato genome (Xu et al., 2011) and the merged potato gene and unigene sequences StNIB_v1 (Ramšak et al., 2014). Analysis of phasing was performed in 21- and 24-nt intervals. To detect *PHAS* loci with a significant degree of phasing, very strict criteria were applied to avoid detection of false positives (phasing p-value < 0.0001; number of unique phasiRNAs detected at specific *PHAS* locus ≥ 4, also to avoid detection of repeat-associated siRNAs). Additionally, the phasing p-values were corrected with strict Bonferroni correction and only loci with the p-value < 0.05 were considered as *PHAS* loci.

### sRNA quantification and statistical analysis

Differential expression analysis was performed in R (R Development Core Team, 2011; version 3.2.2), using the limma package (Ritchie et al., 2015). In short, sRNA counts with a baseline expression level of at least two RPM (reads per million of mapped reads) in at least three samples were TMM-normalized (edgeR package; Robinson et al., 2009) and analysed using voom function (Law et al., 2014). To identify differentially expressed sRNAs the empirical Bayes approach was used and the resultant p-values were adjusted using Benjamini and Hochberg’s (FDR) method. Adjusted p-values below 0.05 were considered statistically significant.

### Stem-loop RT-qPCR

Stem-loop RT-qPCR was used to quantify the expression of six target miRNAs in relation to the endogenous control (stu-miR167a-5p.1), which was determined to be the most robustly expressed in a sRNA sequencing dataset of potato plants that were uninfected or infected with a range of viruses (PVY, PLRV, PVS, PVX, PVT) (SRA accession no. SRP083247). TaqMan MicroRNA Assays (Thermo Fisher Scientific) were ordered according to the sRNA-Seq sequence of the selected miRNAs (**Table S1**). Total RNA (1 μg) of the same samples as used for sRNA-Seq was DNase I (Qiagen) treated and reverse transcribed using SuperScript III First-Strand Synthesis System and stem-loop Megaplex primer pool (both Thermo Fisher Scientific) following the manufacturer’s protocol and previously optimized cycling parameters (Varkonyi-Gasic et al., 2007). No template control RT reactions were set to assess potential Megaplex primer pool background. qPCR reactions were performed in 10 μl volume on the LightCycler480 (Roche Diagnostics Ltd., Rotkreuz, Switzerland) in duplicates and two dilutions (8- and 80-fold) per sample using TaqMan Universal Master Mix II, no UNG (Thermo Fisher Scientific) and TaqMan MicroRNA Assays following the manufacturers’ protocols. Additionally, for each miRNA assay, a standard curve was constructed from a serial dilution of the pool of all samples. Raw Cq values were calculated using the second derivative maximum method (Roche Diagnostics Ltd.) and miRNA expression was quantified using a relative standard curve method by normalization to the endogenous control using quantGenius (Baebler et al., 2017) The statistical significance was assessed by Student’s t-test.

### sRNA target prediction

*In silico* identification of potato transcripts targeted by identified sRNAs was carried out using the psRNATarget ((Dai and Zhao, 2011); http://plantgrn.noble.org/psRNATarget/) and StNIB_v1 sequences (Ramšak et al., 2014), following previously proposed stringent parameters (Zhang et al., 2013). Moreover, targets of identified sRNAs were experimentally validated with parallel analysis of RNA ends (PARE) Degradome-Seq. The four degradome libraries (mock Désirée, PVY Désirée, mock NahG-Désirée, PVY NahG-Désirée) were constructed by pooling RNA of the biological replicates and sequenced on the Illumina HiSeq 2500 platform. The data were analyzed at LC Sciences (Houston, TX) with CleaveLand4 ((Addo-Quaye et al., 2009); http://sites.psu.edu/axtell/software/cleaveland4/) using all our experimentally identified sRNAs and the StNIB_v1 sequences allowing for maximum three mismatches. All identified degradation targets were classified into 5 categories as previously described (Addo-Quaye et al., 2009). Only categories with high confidence of cleavage (0,1,2,3) were considered for biological interpretation (see **Dataset S7** for details). Results of miRNA-target (*PHAS* loci) interactions were also used to reveal miRNA triggers of the phasiRNA production. Only 22-nt miRNAs were kept as potential triggers (Chen et al., 2010; Cuperus et al., 2010).

### Regulatory network construction

In order to compare the expression of sRNAs with the expression of their target transcripts we used a microarray gene expression dataset generated from the same samples ((Stare et al., 2015); GEO accession no. GSE58593). All differentially expressed miRNAs and phasiRNAs were analyzed for functional overrepresentation in biological pathways with MapMan software (Usadel et al., 2009) using the ontology adapted for potato (Ramšak et al., 2014; Rotter et al., 2007). All sRNAs and their targets, obtained by *in silico* prediction and Degradome-Seq were integrated with their expression data and used for the construction of regulatory networks in Cytoscape 3.4 (Shannon et al., 2003).

### Identification of *cis*-regulatory elements in promoter regions of *MIR* genes

1000 nt sequences upstream of the predicted *MIR* gene hairpin sequences were extracted as putative miRNA promoter regions (Megraw and Hatzigeorgiou, 2010) and analysed for *cis*-regulatory elements of plant transcription factors using position weight matrices and transcription binding sites in TRANSFAC (Matys et al., 2003); http://www.biobase-international.com/product/transcription-factor-binding-sites) and PlantCARE (Lescot et al., 2002); http://bioinformatics.psb.ugent.be/webtools/plantcare/html/) implemented algorithms.

### Hormonal measurements

Hormone contents were determined by gas chromatography coupled with mass spectrometry (GC-MS) from approx. 100 mg of plant material. 1 ml of 100% methanol (HPLC grade) and a mix of 50 pmol stable isotope-labelled internal standards were added to each sample prior to extraction. The samples were heated (60 °C, 5 min) and then incubated (room temperature, 1 h) with occasional vortexing. The methanolic phase was taken to complete dryness *in vacuo*. The resulting residue was dissolved in methanol (50 μl) to which diethyl ether (200 μl) was added. The samples were sonified (5 min) and centrifuged (5 min, 14,000 *g*). The particle-free supernatant was loaded to aminopropyl solid-phase extraction cartridges (Chromabond NH_2_ shorty 10 mg; Macherey–Nagel GmbH, Düren, Germany). Each cartridge was washed twice with CHCl_3_:2-propanol (2:1, v/v, 250 μl) before the hormone-containing fraction was eluted with acidified diethyl ether (2% acetic acid, v/v, 400 μl). The eluates were transferred into 0.8 ml autosampler vials and again taken to dryness in a gentle stream of nitrogen. Prior to MS assessment, the samples were derivatized with a 20 μl of a mix of l of acetone:methanol (9:1, v/v, 220 μ), diethyl ether (27 μl) and (trimethylsilyl)diazomethane solution (2.0 M in diethyl ether, 3 μl) and letting them rest for 30 min at room temperature. The setting for the GC and the MS were as described previously (Sanz et al., 2014). For the determination of endogenous and stable isotope-labeled methylated acidic plant hormones, respectively, the following ion transitions were recorded: MeSA m/z 152 to m/z 120 and m/z 156 to m/z 124 for [^2^H_4_]-MeSA, retention time 6.75 ± 0.4 min; MeOPDA m/z 238 to m/z 163 and m/z 243 to m/z 168 for [^2^H_5_]-MeOPDA, retention time 10.00 ± 0.4 min; MeJA m/z 224 to m/z 151 and m/z 229 to m/z 154 for [^2^H_5_]-MeJA, retention time 11.27 ± 0.5 min; MeIAA m/z 189 to m/z 130 and m/z 191 to m/z 132 for [^2^H_2_]-MeIAA, retention time 13.34 ± 0.4 min; MeABA m/z 162 to m/z 133 and m/z 168 to m/z 139 for [^2^H_6_]-MeABA, retention time 15.78 ± 0.4 min; and MeGA m/z 136 to m/z 120 and m/z 138 to m/z 122 for [^2^H_2_]-MeGA, retention time 21.67 ± 0.6 min. The amounts of endogenous hormone contents were calculated from the signal ratio of the unlabeled over the stable isotope-containing mass fragment observed in the parallel measurements. Significant changes in hormone concentrations between treatment-genotype groups were determined by ANOVA followed by LSD post hoc analysis (Benjamini Hochberg FDR p-value adjustment, alpha = 0.05) using the Agricolae R package.

### Gene set enrichment analysis

To identify SA regulated genes in cv. Désirée the normalized expression values between mock NahG-Désirée vs. Désirée and PVY-infected NahG-Désirée vs. Désirée samples were compared (calculated from the 3 dpi samples; data of Stare et al. (2015). Gene Set Enrichment Analysis algorithm (GSEA; (Subramanian et al., 2005) was run to perform analysis (false discovery rate corrected q ≤ 0.01) of expression profiles between both genotypes, using MapMan ontology as the source of the gene sets.

### Data deposition and Gene IDs

The sRNA and Degradome-Seq data can be accessed at the NCBI’s Gene expression omnibus (GEO) under accession numbers GSE84851 and GSE84967; (http://www.ncbi.nlm.nih.gov/geo/query/acc.cgi?token=anqhgqimlhihzen&acc=GSE84967) and will be available after acceptance of the manuscript. Supplemental Online Files 1 and 2 are available at https://www.dropbox.com/sh/yt7qenj7svcryux/AADpRomBYQ1jfyA9a9-9OJuUa?dl=0. A full list of gene/protein names used in this manuscript, together with their Gene IDs, short names, Arabidopsis orthologue genes is given in **Table S2**.

## Results

### Novel endogenous sRNAs identified in potato leaves

We identified 245 different previously described miRNAs (including 38 miRNA variants; isomiRs), belonging to 95 miRNA families in control and PVY^NTN^-infected leaves of cv. Désirée using sRNA-Seq (**Figure S1, Dataset S1**). In addition, 141 novel miRNAs were detected, of those 12 were coded by multiple *MIR* loci. Novel miRNA sequences were assigned to 123 novel miRNA families (**Dataset S1 and S2**).

When assessing the miRNA regulatory network, the amplification of silencing through phasiRNA biogenesis was also considered. In total, more than 400 *PHAS* loci were predicted, coding for 1513 phasiRNAs. 248 *PHAS* loci were located on protein-coding regions of genes, with the majority encoding NBS-LRRs (nucleotide binding site-leucine rich repeat proteins) and LRR-RLKs (leucine rich repeat-receptor-like kinases) (**Dataset S3 and S4**). Moreover, we also searched for miRNAs with the ability to trigger phasiRNA production and similarly to previous reports (Li et al., 2011; Shivaprasad et al., 2012), we observed that the vast majority of predicted miRNA triggers belong to miR482, miR6023, miR6024 and miR6027 families (**Dataset S5**).

We further compared miRNA expression profiles of PVY^NTN^-infected versus mock-inoculated leaves in early stages of virus infection (3 dpi, 1 day before detectable viral multiplication; (Baebler et al., 2011; Stare et al., 2015)). In total, 61 unique miRNAs were found to be significantly differentially expressed in early stages of PVY^NTN^ infection (3 dpi) of Désirée plants. Virus infection predominantly caused an increase in miRNAs levels (**Dataset S1**). Additionally, we identified 36 phasiRNAs as differentially regulated 3 dpi, mainly originating from non-coding *PHAS* and *NBS-LRR* loci (**Dataset S4**). To validate the obtained sRNA-Seq results, abundance of six differentially expressed miRNAs was analyzed by stem-loop RT-qPCR. As shown in **Figure S2**, all sRNA-Seq differential expression results were confirmed.

In previous studies, a plethora of potato miRNA/phasiRNAs has been shown to be differentially expressed following pathogen infection. However, the biological relevance of these differences remains largely unknown. To translate the data obtained on sRNA level into changes in physiological processes, we performed *in silico* sRNA target prediction, both at the levels of translational inhibition and target cleavage (**Dataset S6**). Additionally, the predictions of target cleavage were experimentally validated by Degradome-Seq (**Dataset S7**). Based on this information we constructed a potato sRNA regulatory network connecting miRNAs with phasiRNAs and their targets (**Supplementary Online File 1 and 2**). This revealed several already known and many novel connections linking sRNA regulation to the plant immune signaling (see example in **Figure S3**; **Dataset S6 and S7**). Several miRNA-mRNA pairs conserved across plant species, such as miR156-*SPL11*, miR160-*ARF10*, miR172-*AP2* or miR396-*GRF5* (Curaba et al., 2014), were confirmed also in our system (**Dataset S6 and S7**). Our data also showed the miR3 93-mediated cleavage of transcripts encoding members of *TIR/AFB* gene family, receptors implicated in the control of auxin signaling (Figure 1) (Si-Ammour et al., 2011). We also discovered that these transcripts were targets of several *TIR1*-derived phasiRNAs (phasiTIRs) (Figure 1, **Dataset S6 and S7**). Moreover, our *in silico* analysis predicted that the miR393- and phasiTIR-network is targeting downstream transcription factor StARF1 and other phytohormone signaling pathways, such as transcripts involved in ethylene signaling, in jasmonate signaling and in brassinosteroid biosynthesis (Figure 1, **Dataset S6 and S7**). Two of these alternative phasiTIR targets, *StAP2* and *StOPR1*, were also confirmed by degradome sequencing. Interestingly, our analysis also showed that the majority of differentially expressed miRNAs and phasiRNAs target genes are coding for defense-related proteins such as pathogenesis-related (PR) proteins and proteins involved in the biosynthesis of secondary metabolites, transcription factors belonging to AP2/ERF, bHLH, MYB and GRAS family proteins, putative immune receptors (NBS-LRRs, LRR-RLKs) as well as proteins involved in biosynthesis and signaling of different phytohormones (Figure 2A).

**Figure 1.**
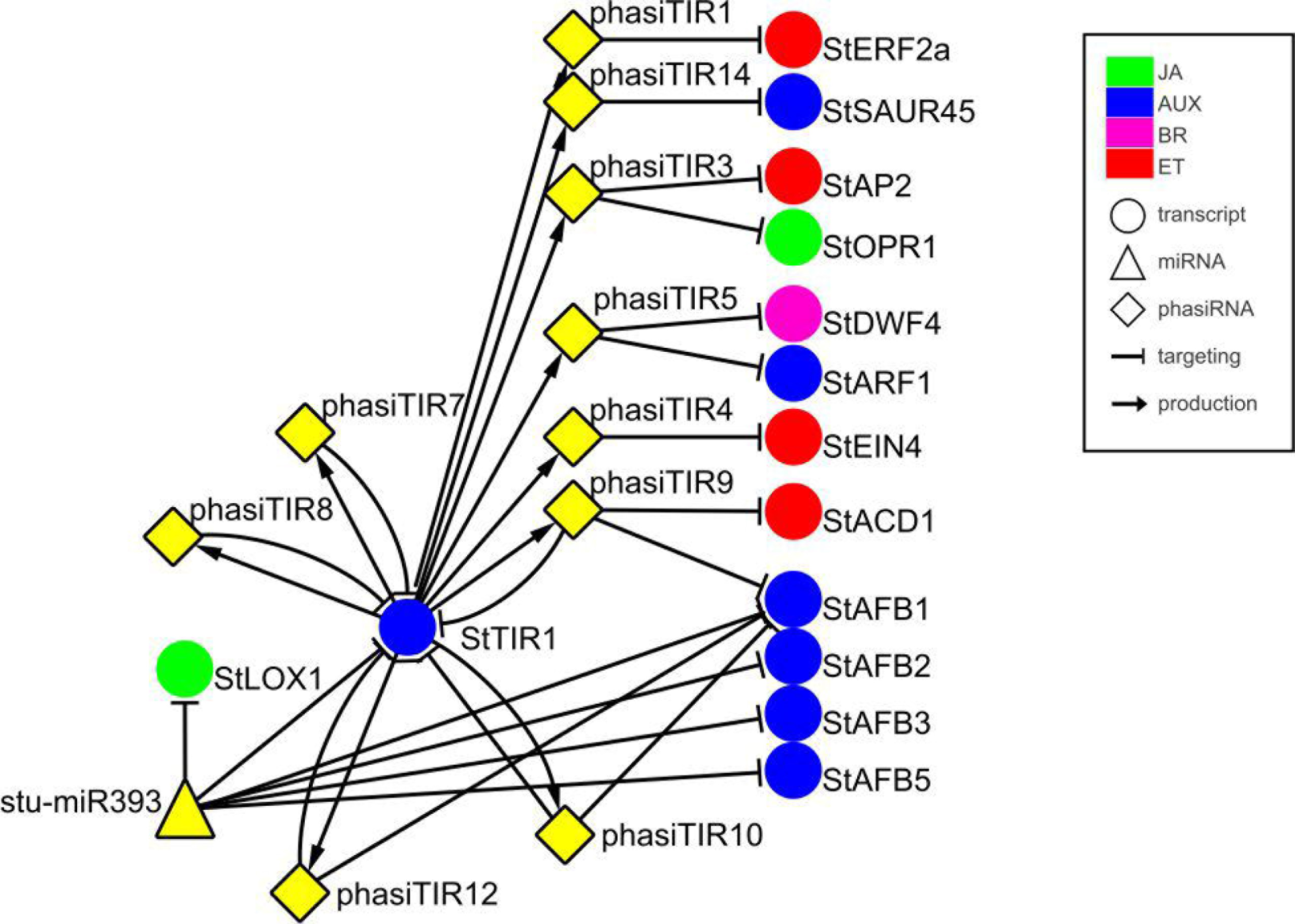
miR393-mediated cleavage of StTIR1 leads to production of phasiTIRs that are predicted to target diverse phytohormone signaling components. Targets of phasiTIRs were predicted *in silico*, cleavage of two of them, *AP2* and *OPR1*, was also confirmed by degradome sequencing (PARE). Node shapes represent classes of sRNAs (triangle – miRNA; diamond – phasiRNA) or transcripts (circle). Node colors indicate components related to different hormone signaling pathways: green – jasmonic acid (JA); blue – auxin (AUX); magenta – brassinosteroid (BR); red – ethylene (ET). Arrows connect sRNAs and targets (blunt-end arrow) or *PHAS* loci and producing phasiRNAs (regular arrow). Node stu-miR393 represents miR393-5p and miR393-5p.1 and node StAFBl represents StAFBl.l, StAFB1.2, and StAFB1.3. For details of the target transcripts/genes see **Table S2**. StTIR1 – Transport inhibitor response 1, StLOX1 – Lipoxygenase 1, StERF2a – Ethylene responsive transcription factor 2a, StSAUR45-Small auxin upregulated RNA 45, StAP2-APETALA2, StOPR1-12-oxophytodienoate (OPDA) reductase, StDWF4 – Dwarf4, StARF1 – Auxin response factor 1, StEIN4 – Ethylene insensitive 4, StACD1 – 1 – aminocyclopropane-1-carboxylic acid deaminase 1, StAFB1/2/3/5 – Auxin F-box 1/2/3/5.

**Figure 2.**
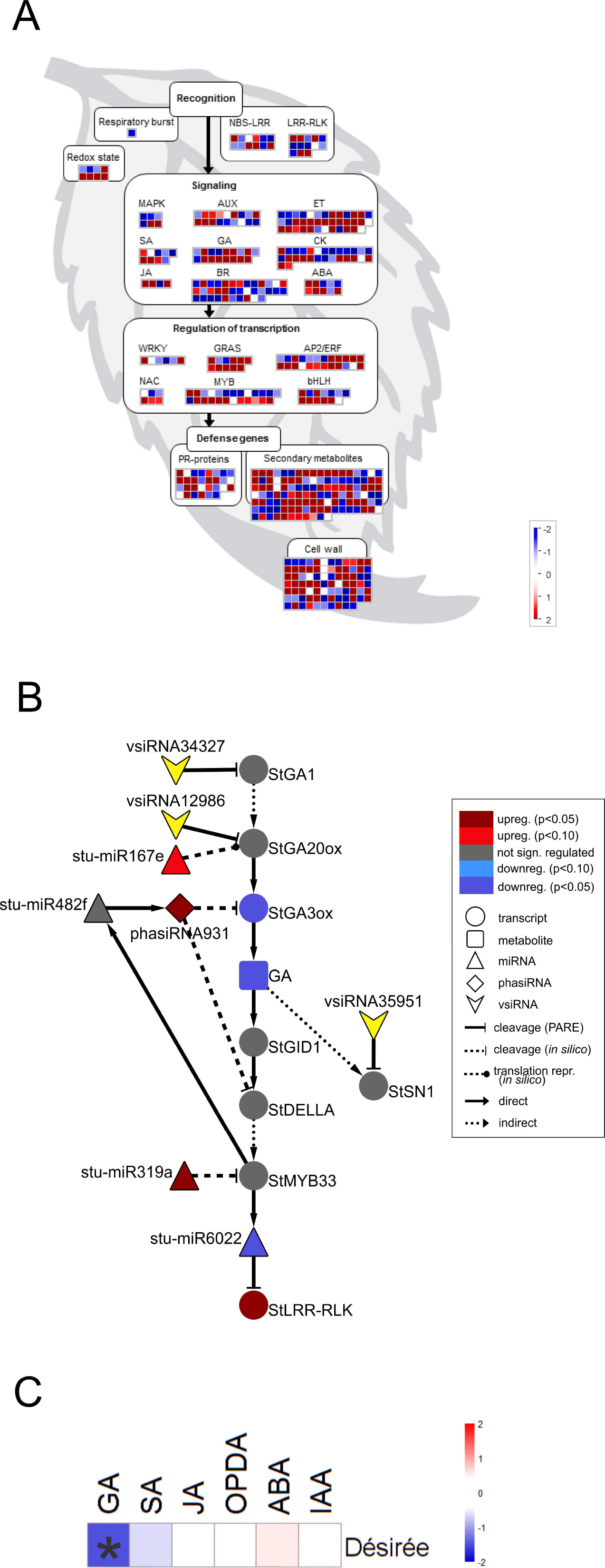
PVY induced changes in sRNA regulatory network are targeting multiple immune and gibberellin signaling components in Désirée at the onset of viral multiplication. (**A**) Visualization of differentially expressed miRNAs/phasiRNAs in PVY^NTN^-infected Désirée according to the function of their predicted targets. Each square represents log_2_ ratios of expression between PVY^NTN^ – and mock-inoculated plants (red – upregulated; blue – downregulated). MapMan ontology bins: respiratory burst (20.1.1), redox state (21.6), MAPK (30.6), SA (17.8), JA (17.7), AUX (17.2), GA (17.6), BR (17.3), ET (17.5), CK (17.4), ABA (17.1), WRKY (27.3.32), NAC (27.3.27), GRAS (27.3.21), MYB (27.3.25), AP2/ERF (27.3.3), bHLH (27.3.6), PR-proteins (20.1.7), secondary metabolites (16), cell wall (10). The NBS-LRR and LRR-RLK bins were custom constructed for this study, based on their harboring domains (obtained from PFAM database; (Finn et al., 2016)). These bins represent differentially expressed miRNAs/phasiRNAs targeting NBS-LRRs or LRR-RLKs. NBS-LRR – nucleotide binding site-leucine-rich repeat protein, LRR-RLK – leucine-rich repeat receptor-like kinase, MAPK – mitogen activated protein kinase, SA – salicylic acid, JA – jasmonic acid, AUX-auxin, GA-gibberellin, BR – brassinosteroid, ET – ethylene, CK – cytokinin, ABA – abscisic acid, PR-pathogenesis-related. (**B**) Network of differentially expressed endogenous sRNAs and vsiRNAs targeting mRNAs of GA biosynthesis and signaling pathways in Désirée 3 days post PVY^NTN^ inoculation. Node shape represent classes of sRNAs (triangle – miRNA; diamond – phasiRNA; arrowhead – vsiRNA), transcripts (circle) or metabolites (rectangle). Statistical significances of expression differences (p-values) and direction of expression change are represented by the node colors (see the legend). Arrows indicate type of interaction (solid-line normal arrow – direct interaction; dashed-line normal arrow – indirect interaction; blunt-end solid arrow-cleavage observed by Degradome-Seq, blunt-end dashed-line arrow – *in silico* predicted cleavage (or translational repression as proposed by Rogers and Chen (2013), dashed-line oval arrow – *in silico* predicted translational repression). Node stu-miR319a represents stu-miR319a-3p and stu-miR319a-3p.2. Node StGA20ox represents StGA20ox, StGA20ox1, StGA20ox3 and StGA20ox4. For details of the target transcripts/genes see **Table S2**. stu-miR167e – stu-miR167e-3p; stu-miR482f – miR482f-3p; stu-miR6022 – miR6022-3p; StGA1 – GA REQUIRING 1 (ent-copalyl diphosphate synthase); StGA20ox – GA20-oxidase; StGA3ox – GA3-oxidase; StGID1C – GA receptor – GA INSENSITIVE DWARF1C hydrolase; StSN1 – Snakin-1; StDELLA – DELLA protein; StLRR – RLK-leucine-rich repeat receptor-like protein kinase. (**C**) Changes in concentrations of a set of plants hormones in cv. Désirée 3 days after PVY^NTN^ infection. sRNA-mediated repression of GA biosynthesis was confirmed by reduced GA_3_ levels in PVY-infected Désirée plants. Colors present as log_2_ ratios of mean concentrations between PVY^NTN^ – and mock-inoculated plants (n=4; red – increased, blue – decreased level). * – indicate statistically significant values (ANOVA; p < 0.05). SA – salicylic acid, JA – jasmonic acid, OPDA – 12-oxophytodienoic acid, ABA – abscisic acid, IAA – indole-3-acetic acid.

### sRNA regulatory network response in tolerant interaction resemble the one in mutualistic symbiosis

Analysis of the differentially expressed miRNAs and phasiRNAs together with the levels of the target transcripts (data published in (Stare et al., 2015)) revealed the presence of many known, as well as novel regulatory cascades involving *NBS-LRR*s. Several *NBS-LRR*s were predicted to be targeted by miR482, miR6024 and miR6027 family members (**Dataset S6 and S7**). Moreover, *NBS-LRRs* are regulated also by phasiRNAs, where most phasiRNAs have multiple *NBS-LRR* targets (**Figure S4**, **Dataset S6 and S7**) due to the shared conserved P-loop or Walker A motif (Shivaprasad et al., 2012). In all of the previous studies, miR482 family members were downregulated following pathogen infection (Ouyang et al., 2014; Shivaprasad et al., 2012; Yang et al., 2015). In our study, however, the one regulated member of the miR482 family (miR482e) targeting *NBS-LRR* transcripts was upregulated following PVY^NTN^ infection (**Dataset S1**), similarly as observed in the establishment of mutualistic symbiosis in soybean roots (Li et al., 2010). Moreover, several miRNAs that were upregulated in response to PVY^NTN^ in cv. Désirée, such as miR164, miR167, miR169, miR171, miR319, miR390 and miR393 have also been reported to regulate nodulation and arbuscular mycorrhizal symbiosis in different legume species (**Dataset S1**; (Devers et al., 2011; Lelandais-Brière et al., 2009; Mao et al., 2013; Yan et al., 2015). In addition to NBS-LRR proteins, LRR-RLKs are also important mediators of immune as well as important triggers of mutualistic symbiosis signaling cascades (Antolin-Llovera et al., 2014; Hohmann et al., 2017). We have predicted a novel miRNA-LRR-RLKs interaction in which miR6022 levels decrease in response to PVY^NTN^ infection in cv. Désirée, which is further linked to upregulation of its predicted target genes encoding LRR-RLKs (**Figure S4A**, **Dataset S6**).

### Several PVY^NTN^-derived siRNAs trigger degradation of host transcripts

The primary plant defence mechanism against invading viruses is RNA silencing involving the production of vsiRNAs. The population of vsiRNAs detected in the infected samples consisted of more than 46 000 unique sequences of 20-24 nt in length (**Dataset S8**). In order to take into account the unlikely possibility that PVY^NTN^ produces its own miRNAs, we first ran the miRNA prediction pipeline on vsiRNAs and the viral genome. However, we found no sequence that would fulfil the criteria for a viral miRNA. Subsequently, we searched for potential host transcripts targeted by vsiRNAs in our experimentally validated target degradation dataset (**Dataset S9**). We found that vsiRNAs are indeed able to target multiple potato transcripts, among them mRNAs coding for immune receptor proteins, various transcription factors and proteins involved in hormonal signaling pathways (**Dataset S9**). For example, several vsiRNAs were detected with the confirmed ability to guide the cleavage of transcripts involved in auxin signaling, transcripts encoding IAA-amino acid hydrolases (StILR1 and StIAR3), Aux/IAA transcriptional repressors (StIAAs) and the transcription factor StARF2.

### sRNA-mediated downregulation of GA biosynthesis genes is reflected in lower GA_3_ levels

Interestingly, we found that GA biosynthesis and downstream signaling are targeted by a sRNA-mediated regulatory network and that the changes in sRNA levels following PVY^NTN^ infection are reflected also in the changes of their target transcripts levels (Figure 2B, **Dataset S6**). GA20-oxidase (GA20ox) and GA3-oxidase (GA3ox) are enzymes that catalyze the last steps in the formation of bioactive GAs (Yamaguchi, 2008). In Désirée plants miR167e was predicted to cause translational repression of the *StGA20ox* transcript (Figure 2B). An additional layer of GA biosynthesis regulation is represented by the increased production of phasiRNA931, which promotes cleavage of the *StGA3ox* transcript (Figure 2B). The transcriptomics results support these interactions as the targeted transcripts are significantly downregulated in Désirée upon PVY^NTN^ infection (Figure 2B, **Dataset S6**). Additionally, vsiRNAs were found to target transcripts encoding two enzymes involved in GA biosynthesis *StGA1 and StGA20ox* (Figure 2B, **Dataset S9**). One also has to note that all of the miRNAs/phasiRNAs discovered to be involved in sRNA-GA biosynthesis regulation have so far not been identified in Arabidopsis and among them only miR167e was also discovered in tomato (Griffiths-Jones et al., 2006; Zhang et al., 2014). sRNAs can target downstream GA signaling in the potato-PVY interaction on multiple levels. Four miR319 family members, close relatives of miR159 family (Palatnik et al., 2007), were predicted to cleave the transcript encoding StMYB33, an orthologue of GAMYB transcription factor involved in GA signal transduction (Millar and Gubler, 2005). Moreover, phasiRNA931 can cleave a potato orthologue of a DELLA protein, a GA-signaling repressor (Figure 2B, **Dataset S6**).

Such interconnectedness between plant defense-related miRNA/phasiRNA network and GA biosynthesis/signaling has not been previously identified and may represent a link between defensive and developmental signaling. Thus, we decided to functionally evaluate these results by measuring the concentrations of a set of plant hormones. As predicted by the sRNA-target transcript analyses, we detected a reduced level of GA3 in PVY^NTN^ infected Désirée plants (Figure 2C). The levels of SA, jasmonic acid (JA), the JA precursor 12-oxophytodienoic acid (OPDA), abscisic acid (ABA) and indole-3-acetic acid (IAA) remained unchanged at 3 dpi (Figure 2C, **Dataset S10**). To inspect if GA deficiency has any impact on plant growth, plant height was monitored till 21 dpi. No differences were observed between all four studied groups of plants.

### *In silico* prediction showed interconnection between miRNA regulatory network and transcriptional regulation

Given the critical role of miRNAs in gene regulation, *cis*-regulatory elements of differentially expressed *MIR* genes involved in *R*-gene regulation and GA signaling, *MIR6022*, *MIR319a* and *MIR167e* were investigated together with *MIR482f* and *MIR482c* genes encoding miRNA triggers of phasiRNA931 production. Interestingly, GAMYB binding sites were detected in the promoters of the *MIR6022* and *MIR482f* genes (**Dataset S11**). Moreover, these genes harbor WRKY8/28/48 binding sites, while *MIR319a* and *MIR482c* harbor a general WRKY binding W-box regulatory element. Additionally, *cis*-acting elements involved in SA and JA responsiveness were identified in the promoter of the *MIR482f* gene (**Dataset S11**). In all these four analyzed promoter sequences NAC transcription factor binding sites were detected. The promoter of *MIR167e* is only partially assembled in the current version of the potato genome; thus, the promoter analysis was performed only for the first 80 nt upstream of the predicted hairpin precursor sequence (**Dataset S11**).

### SA depletion attenuates sRNA response following PVY^NTN^ infection

As the link between repression of GA signaling and disease symptoms severity was already established in Arabidopsis (Du et al., 2014) we have further investigated the activity of discovered sRNA-GA circuit in susceptible potato-PVY^NTN^ interaction. We have shown previously that SA depletion breaks the equilibrium between disease and tolerance. In NahG-Désirée plants, the pronounced disease symptoms appeared both on the inoculated and systemic leaves (Baebler et al., 2011). Furthermore, the viral multiplication was detected one day earlier than in non-transgenic Désirée plants (at 4 dpi), while the final concentrations of the virus were not significantly higher (Baebler et al., 2011). Here, we performed the analysis of sRNA response as well as the measurements of hormonal concentrations in the interaction of this susceptible genotype with the virus. Interestingly, we found that the overall response of sRNAs was attenuated in NahG-Désirée at 3 dpi. In NahG-Désirée only 28 miRNAs were differentially expressed, with the majority showing a lower degree of induction than in Désirée plants (Figure 3 **and 4A, Dataset S1**).

**Figure 3.**
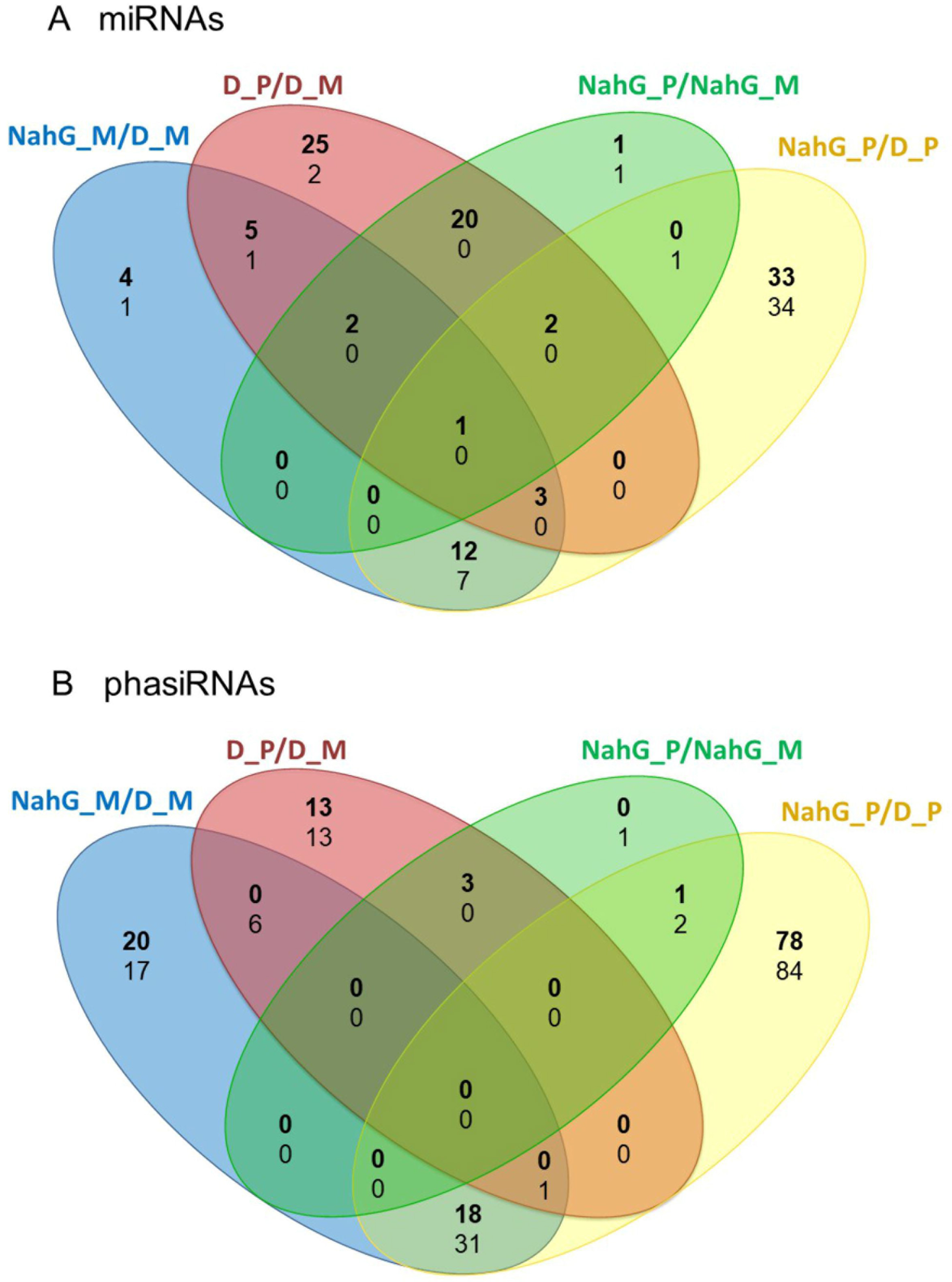
Numbers of unique and common differentially expressed miRNAs and phasiRNAs 3 days post PVY^NTN^ inoculation in comparison of SA-deficient and non-transgenic Désirée plants. Venn diagrams show the number of differentially expressed (FDR corrected p-value < 0.05) (**A**) miRNAs and (**B**) phasiRNAs in mock- or PVY^NTN^-inoculated potato leaves of cv. Désirée and NahG-Désirée. Upregulated miRNAs/phasiRNAs are shown in boldand downregulated in normal text. D – Désirée, NahG – NahG-Désirée, M – mock, P – PVY^NTN^.

Inspecting specifically the discovered link between sRNAs regulation, GA signaling and immune signaling in the sRNA and transcriptional datasets we observed that the responses of miR167e and miR319a were diminished in NahG-Désirée plants (Figure 4B, **Dataset S1**). Interestingly, although the phasiRNA931 was also upregulated in NahG-Désirée plants, albeit to a lower extent, this was not translated into downregulation at the target transcript level (Figure 4B, **Dataset S6**). Adding to the significance of this finding, in NahG-Désirée plants, the level of bioactive GA was not significantly different in infected leaves (Figure 4C). Furthermore, the relieved silencing of LRR-RLKs by miR6022 that is predicted to modulate the immune response and which is also linked with GAMYBs was absent in NahG-Désirée plants (**Figure S4B**). We also inspected sRNA-mediated responses which resembled responses in mutualistic symbioses in tolerant plants of Dèsirèe and found that miR482e was also upregulated in NahG plants, while regulation of miR164, miR167, miR169, miR171, miR319, miR390 and miR393, was diminished (**Dataset S1**).

**Figure 4.**
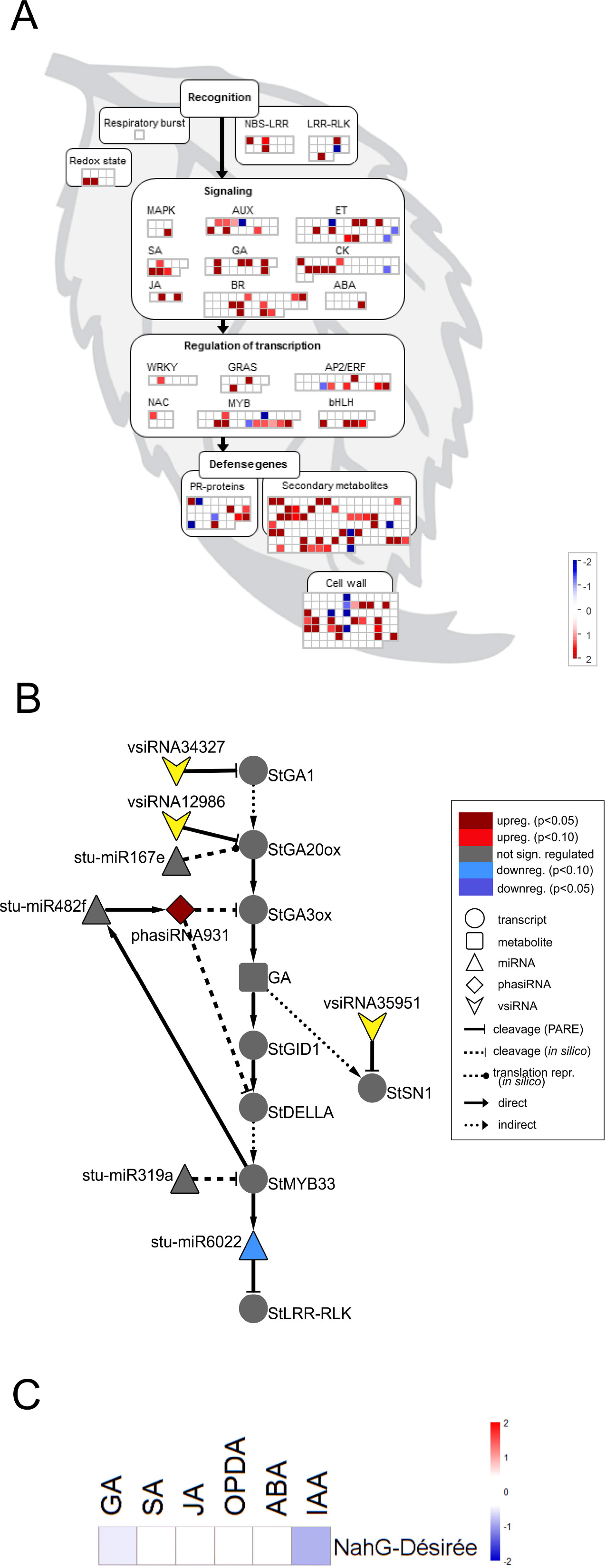
sRNA response is attenuated in susceptible SA depleted plants following PVY^NTN^ infection. (**A**) Visualization of differentially expressed miRNAs/phasiRNAs in PVY^NTN^-infected NahG-Désirée according to the function of their targets. Each square represents log2 ratios of expression between PVY^NTN^ – and mock-inoculated plants (red – upregulated; blue – downregulated). (**B**) Network of endogenous sRNAs and vsiRNAs targeting mRNAs of GA biosynthesis and signaling pathways in NahG-Désirée 3 days post PVY^NTN^ inoculation. (**C**) Concentrations of a set of plants hormones in NahG-Désirée 3 days after PVY^NTN^ infection. The levels of all analyzed hormones remained unchanged in NahG plants following PVY^NTN^ infection. Colors present as log_2_ ratios of mean concentrations between PVY^NTN^ – and mock-inoculated plants (n=4; red – increased, blue – decreased level). * – indicate statistically significant values (ANOVA; p < 0.05). For abbreviations and other details of the scheme, see the caption of Figure 2.

To evaluate the direct effect of the SA deficiency in NahG plants, independently of the viral infection, we also compared the sRNA expression profiles in mock-inoculated leaves of non-transgenic and SA-depleted Désirée (Figure 3). We found 36 miRNAs regulated by SA. It seems that SA in the normal growing conditions generally cause the reduction in the levels of miRNAs as the majority (27) of miRNAs were detected at significantly higher levels in NahG-Désirée plants (**Dataset S1**). When we similarly compared the transcript expression profiles of the two genotypes we noticed that the most strongly induced by SA signaling are notably different WRKY transcription factors (Table 1, **Dataset S12 and S13**), among them orthologues of Arabidopsis *WRKY70*, which was already shown to be positively regulated by SA (Li et al., 2004). As *MIR319a*, *MIR482c* and *MIR482f* promoters harbor WRKY transcription factors binding sites (**Dataset S11**), we discovered a potential direct link between SA signaling and miRNA regulatory network in potato. This link was experimentally confirmed by differential response of these three miRNAs to PVY infection in non-transgenic versus SA-depleted genotype. miR319a was only induced following PVY infection in non-transgenic plants, supporting the hypothesis that it requires WRKY transcription factor for this response. On the other hand, miR482c and miR482f accumulation only differs in SA-depleted plants indicating more complex regulation indicated also by prediction of additional SA responsive elements in their promoter regions (Figure 5, **Dataset S1 and S11**).

**Table 1.**
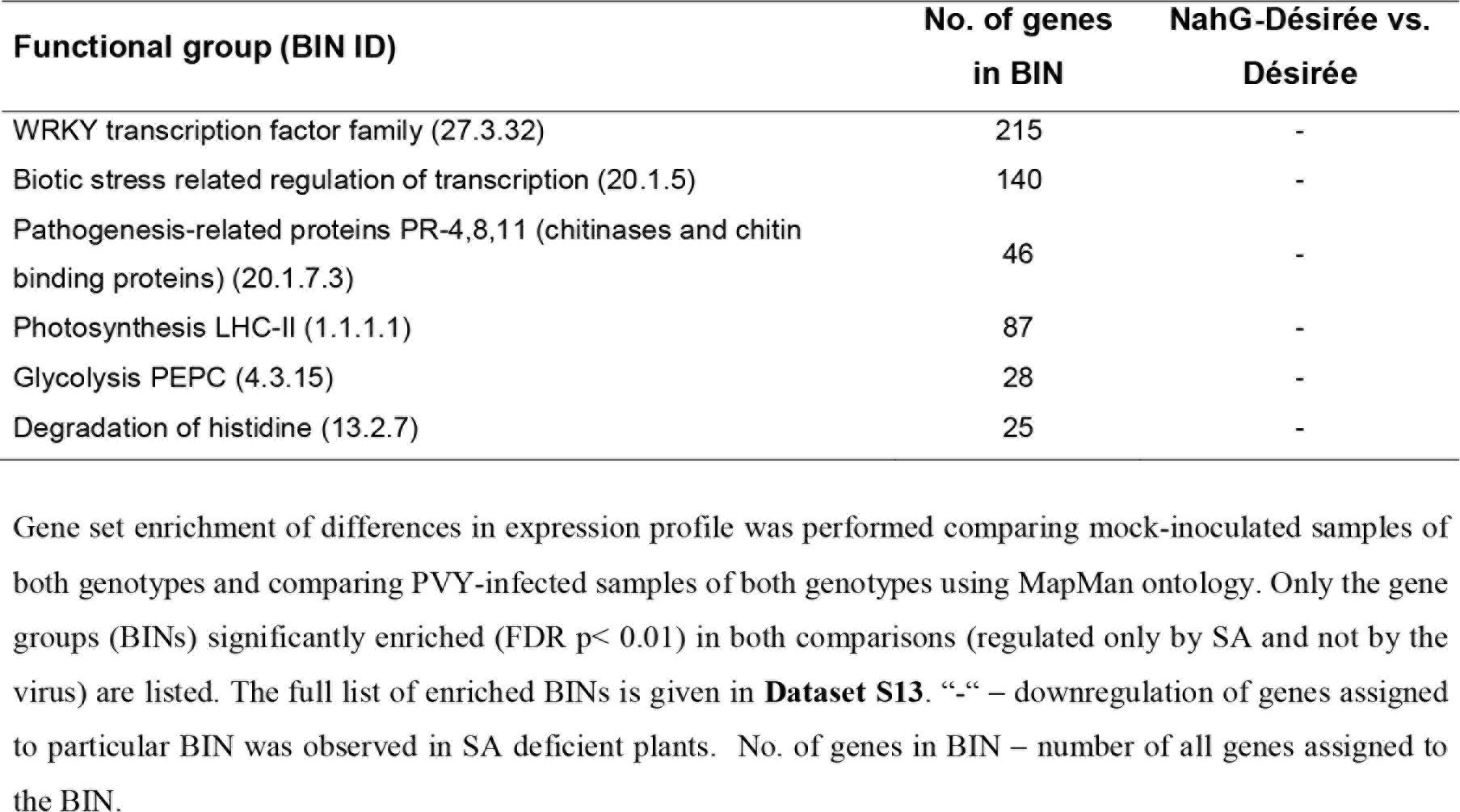
SA-dependent transcriptional responses of potato leaves in cv. Désirée.

| Functional group (BIN ID) | No. of genes<br>in BIN | NahG-Désirée vs.<br>Désirée |
| --- | --- | --- |
| WRKY transcription factor family (27.3.32) | 215 | - |
| Biotic stress related regulation of transcription (20.1.5) | 140 | - |
| Pathogenesis-related proteins PR-4,8,11 (chitinases and chitin binding proteins) (20.1.7.3) | 46 | - |
| Photosynthesis LHC-II (1.1.1.1) | 87 | - |
| Glycolysis PEPC (4.3.15) | 28 | - |
| Degradation of histidine (13.2.7) | 25 | - |

**Figure 5.**
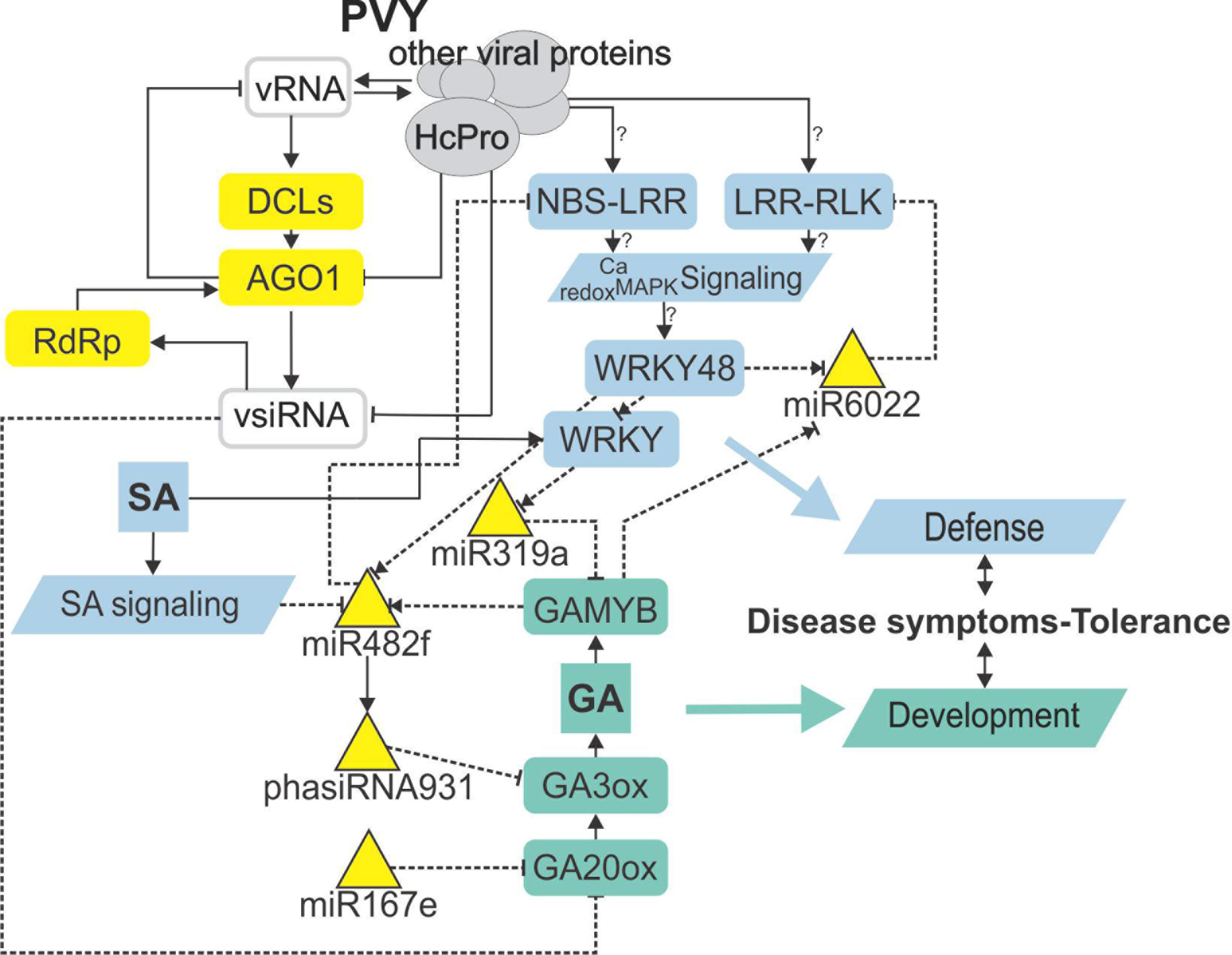
sRNA regulatory network is intertwined with immunity- and gibberellin-related signaling mediating trade-offs in development and defense. Node color denotes component type/function (grey – virus-derived; yellow – RNA silencing; blue – immune response; green – plant development). Lines represent different types of interaction (solid line – protein level; dashed – transcriptional/post-transcriptional level). Normal arrow – activation, blunt-end arrow – inhibition, combination of normal arrow and blunt-end arrow – unknown mechanism of action. ? – inferred from experiments performed in other species. vRNA – viral RNA; vsiRNA – virus-derived siRNA; HcPro – helper component-proteinase; DCL – DICER-like protein; AGO1 – Argonaute 1; RdRp – RNA-dependent RNA polymerase; NBS-LRR – nucleotide binding site-leucine-rich repeat protein; LRR-RLK – leucine-rich repeat receptor-like kinase; Ca – calcium; MAPK – mitogen activated protein kinase; SA – salicylic acid; GAMYB – GA-induced MYB-like protein; GA – gibberellin; GA20ox – GA20-oxidase; GA3ox – GA3-oxidase.

Gene set enrichment of differences in expression profile was performed comparing mock-inoculated samples of both genotypes and comparing PVY-infected samples of both genotypes using MapMan ontology. Only the gene groups (BINs) significantly enriched (FDR p< 0.01) in both comparisons (regulated only by SA and not by the virus) are listed. The full list of enriched BINs is given in **Dataset S13**. “−“ – downregulation of genes assigned to particular BIN was observed in SA deficient plants. No. of genes in BIN – number of all genesassigned to the BIN.

## Discussion

We hypothesized that fine-tuned regulation of subsets of genes involved in defensive signaling can interfere with developmental signaling, which could explain decreased symptom severity in plants expressing tolerance to virus infection. The sRNAs have proven to be an important level for precise regulation of several developmental processes (Fouracre and Poethig, 2016). We here show that the integration of sRNA network and transcriptional regulation is also crucial in the entanglement of immune responses and developmental processes.

Integration of the sRNA expression and expression profiles of their targets confirmed many known, but also revealed some novel regulatory circuits associated with immunity regulation and hormone signaling (Figure 1 **and 2**, **Figure S3 and S4**, **Dataset S6 and S7**). When plants are exposed to pathogens, NBS-LRRs and LRR-RLKs are the key players in sensing and transducing stress signals (Antolin-Llovera et al., 2014; Jones and Dangl, 2006). Viral suppressors of silencing can release the tight control of *R*-gene silencing by sRNAs and activate immune responses in plants (Li et al., 2011; Shivaprasad et al., 2012). Our study investigated regulatory processes occurring early after infection, before virus concentration significantly increased thus the effects we detected were not caused by extensive HCPro or any other viral protein accumulation. Even so, we have detected diverse regulation of NBS-LRRs and LRR-RLKs and their targeting miRNAs as expected according to their specialized roles (**Figure S4**). On the other hand, similarity of response between mutualistic symbiosis in legumes and tolerance in potato was shown by the miR6022-relieved silencing of LRR-RLKs as well as by profiles of several other miRNAs (**Figure S4A**, **Dataset S1**; (Arikit et al., 2014; Devers et al., 2011; Lelandais-Brière et al., 2009; Mao et al., 2013; Yan et al., 2015)). This suggests a similar sRNA network modulation of immune response and physiology occurs in both mutualistic and disease tolerant (commensalistic) interactions. In tolerance, plants may adopt some processes resembling mutualistic symbiosis to control plant response and minimize severe disease symptoms allowing non-hindered development of the plant and at the same time multiplication of the virus.

Phytohormones modulate plant defence responses against various biotic and abiotic stresses as well as plant growth and development (Huot et al., 2014). Till now, several miRNAs were confirmed to participate in this complex network, mainly in connection to repression of auxin signaling (Navarro et al., 2005). A similarly complex miR393/miR396/phasiTIR auxin signaling network was predicted in this study, yet showing links also to other hormonal signaling pathways (Figure 1, **Figure S3**).

Most notable is, however, the novel link between sRNA regulatory network and GA biosynthesis and signaling. Biotic stress was shown to repress GA signaling pathways (Wang et al., 2007). Here, we show that GA biosynthesis and signaling are post-transcriptionally regulated via multiple miRNAs, phasiRNAs as well as vsiRNAs in potato leaves following infection with PVY^NTN^ (Figure 2B). The effect of this regulatory circuit was confirmed by reduced bioactive GA level in the tolerant Désirée plants (Figure 2C). This reduction was however not reflected in decreased plant growth and was thus most probably transient and localized in nature (Karasov et al., 2017). In other plant species, GA biosynthesis was shown to be indirectly targeted by miRNAs regulating the activity of the corresponding transcription factors (miR319-TCP14-GA2ox/GA20ox; miR393-GRF2-GA3ox/GA20ox) (Curaba et al., 2014), while direct interactions were to our knowledge not yet reported. Also of note, the sRNAs regulating GA biosynthesis identified here (miR482f/phasiRNA931 and miR167e) were not identified in Arabidopsis and seem to be Solanaceae specific. The functional relation between lower activity of GA signaling was already directly confirmed to be related to disease severity in three different experimental systems. Arabidopsis *GAMYB* double knockout showed ameliorated symptoms when infected with *Cucumber mosaic virus* (CMV) (Du et al., 2014), similarly was shown in rice in interaction with bacteria (*Xanthomonas oryzae* pv. *oyrzae*) and fungi (*Magnaporthe oryzae*) using knockout in GA deactivating enzyme (Yang et al., 2008). Also in line with this, decrease in GA levels and increase in DELLA protein concentrations was shown to trigger components of rhizobial and mycorrhizal signaling (Calvo et al., 2004; Floss et al., 2013; Fonouni-Farde et al., 2016; Ghachtouli et al., 1996; Jin et al., 2016), showing yet another similarity between tolerant response of potato to viral infection and response of plants in mutualistic symbiosis.

With the discovery of GAMYB binding sites in the *MIR482f* and *MIR6022* promoters (**Dataset S11**), we predicted the circuit in sRNA-GA signaling and additionally a link between GA signaling and *R*-gene expression. The complexity of regulatory responses observed (Figure 5) is in line with the systems biology paradigm that interaction of multiple components and not a single component within a cell leads to much of biological function (Westerhoff et al., 2015). Although the reductionist approach is powerful in building logically simple hypotheses and devising ways to test them, it is very difficult to reconstitute the function for a whole biological system based solely on that as the behaviour of the system may depend heavily on complex interactions within the system (Hillmer et al., 2017). Thus, we have adopted a systems level confirmation of our hypothesis that sRNA-GA-immune signaling interaction is important for the establishment of tolerant interaction. Previously, we had demonstrated that SA regulates plant responses to virus infection; not only by delaying viral multiplication, but also by controling disease symptom severity, most probably via its effects on host primary metabolism (Baebler et al., 2011). In this study, we have shown that response of sRNA regulatory networks controling potential immune receptors and hormonal signaling is strongly attenuated in the NahG-transgenic plants in the early stage of viral infection (3 dpi; Figure 3 **and** 4) linking the sRNAs regulatory network, immune signaling and symptoms development. The molecular mechanisms of the link between SA signaling and sRNA network are also complex. SA has been shown to induce *RNA-dependent RNA polymerase 1* expression, which is crucial for the maintenance of basal resistance to several RNA viruses (Carr et al., 2010; Yu et al., 2003) but none of the silencing mechanism related enzymes is regulated in SA deficient NahG-Désirée plants (**Dataset S12**; (Stare et al., 2015)). We have here predicted and experimentally confirmed SA-directed transcriptional regulation of *MIR482f*, the miRNA linking the GA signaling circuit and R-gene expression (Figure 5, **Dataset S1, S6 and S10**), which could be an additional link between SA signaling and symptoms development in potato-PVY interaction. A direct link could be the WRKY transcription factors that are under the positive control of SA (Table 1, **Dataset S12 and S13**) and were predicted to regulate promoters of all three miRNAs involved in the sRNA-GA circuit (Figure 5, **Dataset S11**).

The outcome of plant-pathogen interactions depends on the delicate balance between plant immune signaling network and its interaction with the pathogen. Here, we focused on the roles of sRNA networks in establishment of the tolerance to PVY infection. We showed that the sRNA regulatory network links immune and developmental signaling in potato through newly discovered sRNA-GA circuit. In tolerance, virus infection perturbs sRNA network resulting in downregulation of GA-mediated signaling, as well as modulation of *R*-gene transcript levels; this results in ameliorated disease symptoms. Supporting this, the responses observed for individual miRNAs are similar as observed in establishment of mutualistic symbioses. It is thus plausible that a similar modulation of plant responses occurs in both mutualistic symbiosis and tolerance. This is in line with growing evidence showing that viruses, like other symbionts, lie on a continuum between antagonistic and mutualistic relationships (Kamitani et al., 2016; Roossinck, 2015).

## Author Contributions

MK, MP, DD, ŠB, JK, JZ, and KG conceived and designed the research; MK, DD, ŠB, and SP performed the experimental work; MK, MP, ŽR, and KG analyzed the data; and MK, ŠB, SP and KG wrote the manuscript. All of the authors have read and approved the final version of the manuscript.

## Acknowledgments

The authors would like to acknowledge dr. Tjaša Stare for providing plant material, dr. Sabine Rosahl for providing NahG-Désirée potato plants. Katja Stare and Tjaša Lukan for technical support, Maja Zagorščak for providing the script to determine the location of *MIR* loci in the potato genome, dr. Denis Kutnjak for the help with PVY^NTN^ genome assembly and dr. John Carr and dr. Anna Coll Rius for critical reading of the manuscript and fruitful discussions.

